# Learning Equivariant Object Recognition and its Reverse Application to Imagery

**DOI:** 10.1101/2023.05.20.541553

**Authors:** Florentine Klepel, Rainer Goebel

## Abstract

To investigate the relationship of perception and imagery, we model the visual ventral stream with an encoder and decoder part with the help of capsule networks. The proposed network consists of V1 and V2 from CorNet-Z, as well as the Capsule Network architecture with the routing by agreement algorithm for V4 and IT. The decoder reverses this architecture to model the feedback activation patterns of the visual ventral stream. The model was trained using EMNIST (letters H, S, C, T). Resulting classification performance was high with good generalization performance to different sizes, positions, and rotations. Contextual information was used for occluded stimuli in the feedback path for reconstructions resulting in high classification performance. Additionally, a pre-trained network was used to reconstruct remapped fMRI activation patterns from higher visual areas. Reconstructions of single-trial imagery data showed significant correlations to physical letter stimuli. The fMRI activation patterns of V1 and V2 and their reconstructions with population receptive field mapping and an autoencoder were related to activation patterns of the network to test biological plausibility. Representational Similarity Analysis and spatial correlations indicated an overlap of information content between the capsule network and the fMRI activations. Due to the capsule networks’ high generalization performance and the implemented feedback connections, the proposed network is a promising approach to improve current modelling efforts of perception and imagery. Further research is needed to compare the presented network to established networks that model the visual ventral stream.

## Introduction

### Deep Neural Networks and Object Recognition

The ventral visual stream is assumed to be relied upon during object recognition in primates (Poggio & Anselmi, 2016). Artificial neural networks (ANNs) modelling the corresponding brain areas are utilized to understand biological vision (Heinke et al., 2022). Even though the human brain performs object classification effortlessly, this level of performance has proven to be difficult to reproduce with artificial systems that are neurobiologically plausible (Poggio & Anselmi, 2016). ANNs showed similar classification performance as the macaque visual system but otherwise could not explain the hierarchical architecture in the cortex (Kubilius et al., 2018). The high complexity of having 100 layers compared to 4 to 8 areas in the visual stream and the lack of recurrent connections are named as main shortcomings of state-of-the-art neural networks to model brain activation during image classification.

Most of these ANNs extract abstract visual properties of the big image data sets used for training various layers (Zhao et al., 2017). These computations are highly susceptible to variations in the input characteristics, such as translations, scaling, and rotations. Scale and position invariance can be achieved by scanning images at different positions and scales (Serre et al., 2007). But when unknown transformations were introduced in the testing data, the performance decreased strongly. Suggested ways to overcome this lack of invariance representation are data augmentation or feature averaging (Lyle et al., 2020). Zhao et al. (2017) used marginalization over transformation parameters to increase insusceptibility to transformations. Other computational models such as neocognitron, and HMAX achieve some level of invariance by alternating feature detector layers with layers that perform local pooling and subsampling of the feature maps (Goodfellow et al., 2009). These ANNs implement restricted forms of invariance but often lack neurobiological validity since their main goal is to reach fast and accurate object recognition which is why they are less useful in explaining neural activity during object recognition in humans.

Serre et al. (2007) argue that the human visual system can compute representations of images that are invariant to the most common image transformations. They hypothesize that the main computational goal of the ventral stream during development might be to learn how objects transform. This gives the visual system the ability to compute versions of new images that are automatically invariant to the same transformations. That indicates that the ventral stream neuronal activity is shaped by invariance representations (Serre et al., 2007). Relevantly, current ANNs still need a large amount of training data which is unrealistic regarding human learning processes. That might indicate that they do not exploit invariance or the relation between features to a large enough extent yet (Bowers et al., 2022).

Convolutional Neural Networks (CNNs), a subcategory of ANNs, reach high performance on automatic visual recognition tasks. They contain successive layers of convolution and pooling with the lower areas explaining the activity in V1, V2, and V3 while activity in higher-level areas in the ventral visual stream are better explained by higher layers (Ramakrishnan et al., 2015). Even though models explaining V1, V2, V4, and IT (occipital to inferior temporal cortex) data were built, their performance is less robust regarding image degradations such as contrast reduction or additive noise, compared to human performance (Geirhos et al., 2017). CNNs and ANNs in general lack performance when dealing with flexible and highly variable objects, and when background noise is high (Poggio & Anselmi, 2016).

Peters et al. (2012) hypothesize that implementing algorithms derived from neurobiological findings might increase the accuracy of computational models. Research made clear that V1 activity of the visual pathway is well described by Gabor-like edge detectors. That is why the same object can elicit very different neural activation patterns in this area when presented in different locations or at different rotation levels (Peters et al., 2012) but invariant representations emerge in final stages of the ventral stream. Even though it was shown that an explicit gradient for feature complexity exists in the ventral pathway (Güçlü & van Gerven, 2015), it is unclear which specific function is fulfilled by which area in the ventral stream. Multiple transformations are likely performed at the intermediate stages (Peters et al., 2012). When investigating the higher-level activity of this stream, Kar et al. (2019) found improved predictions of late IT neural unit responses in shallow recurrent CNNs compared to feedforward-only deep CNNs. They argue that recurrent circuits are crucial for rapid object identification since primates were able to outperform feedforward-only deep CNNs for classifying challenging images that required additional recurrent processing beyond the initial feedforward IT response. This finding illustrates that implementing neurobiological findings into ANNs might in turn lead to improved theories about the neurobiology of the human brain (Peters et al., 2012).

### Capsule Networks

Sabour and colleagues (2017) proposed capsule networks (CapsNets) which might be able to overcome some of the already named shortcomings of conventional ANNs. Their network does not rely on pooling for information reduction and invariant object classification. Instead, the relative position of features is used to detect samples of categories. They showed that a group of units in the hierarchically uppermost layer can represent both the probability and the instantiation parameters of detected entities so that a detailed reconstruction from solely the output vector is possible. The probabilities are invariant representations of the stimuli whereas the instantiation parameters reflect equivariance. Equivariance refers to the same sample eliciting different activation patterns depending on physical features of a specific stimulus, such as position or size (Qiao et al., 2018). CapsNets incorporate an equivariant representation whereas most CNNs only represent invariance at their highest-up layer. It has been shown that viewpoint changes by two pixels in each direction lead to simple linear effects on the pose parameters representing the relationship between an object part and the whole (Mazzia et al., 2021). CapsNets classification accuracy to unseen affine transformations was higher than with a traditional convolutional model (Sabour et al., 2017). Overall, CapsNets have a high intrinsic generalization capability because they rely on object-part relationships instead of feature filters.

The CapsNet includes a routing-by-agreement algorithm which computes and adjusts predictions depending on lower-level capsule predictions of parent-capsule outputs and the coefficient between their actual output and the parent capsule output (Pucci et al., 2021). With the help of this iterative routing process, parts are assigned to wholes. This mechanism also made it possible for the network to recognize multiple objects in an image even when the objects overlap.

Capsule networks can fulfil similar tasks as conventional ANNs. They have already been used for diverse tasks, such as detecting fake images and videos, recognizing human movements, learning time information from spatial information, encoding facial actions, classifying hyperspectral images, finding relationships during natural language processing, classifying emotions, action video segmentation, as well as predicting Alzheimer disease, classifying apoptosis, identifying sign language, and classifying brain tumour types (Huang & Zhou, 2020). Additionally, they were also used for understanding the human brain. Qiao et al. (2018) demonstrated that the reconstruction of capsules fed and trained on fMRI perceptual data is highly accurate. This indicates that there is a common space between these highest level capsule activations and the human brain activations during number perception. However, only categories “6” and “9” of the MNIST digit data set were taken into account.

### Perception and Imagery

Activation patterns during visual mental imagery in early visual areas have been shown to resemble those elicited when sensory stimuli were presented to the participant. The pathway that is used to project perceptual activity upwards in the ventral stream, is assumed to project backwards during mental imagery (Pearson, 2019). Senden et al. (2019) investigated mental imagery and perception of letters in healthy participants. With 7T functional magnetic resonance imaging (fMRI), they investigated whether imagined letter shapes can be reconstructed from V1, V2, and/or V3 activity. Participants were instructed to imagine shapes of the letters H, S, C, and T. Population receptive field (pRF) mapping and an autoencoder were used to transform, augment, and classify the data for the analysis. They showed that the geometric profile of the letter was preserved and could be decoded from early visual areas during imagery. The same early visual areas were relevant to decoding the letter shape during perception and imagery.

It is assumed that top-down feedback processes from higher-level areas are responsible for projecting activity down to early visual areas during mental imagery (Pearson, 2019). Similar activations during perception and imagery were seen in occipital, parietal, and frontal brain areas with an increased overlap in higher-level visual areas (Dijkstra et al., 2019). Dijkstra, et al. (2020) illustrated the feedback imagery process with the help of magnetencephalography (MEG) and a retro-cue task. Participants were presented with two images consecutively, and a cue indicating which one to imagine. With the help of this paradigm, they demonstrated that the information flow during perception is mainly feedforward from lower-level areas to higher-level areas but also included alternating feedback processes. In contrast, perception processes are reactivated in reverse order and mainly top-down feedback processing is involved in imagery. Each feedforward connection in the visual pathway is matched by a reciprocal feedback connection from higher-to-lower-level areas (Gilbert & Li, 2013). Neurons are able to adapt their functioning in a state-dependent manner. They are assumed to constitute processors that adapt their behavior depending on feedback from higher-order cortical areas (Gilbert & Li, 2013). Conventional ANNs mainly contain feedforward connections even though the number of feedback connections is outnumbering the number of feedforward connections in early visual areas (Vetter et al., 2014). Feedback and feedforward weights need to be separated to overcome the unrealistic symmetry in connections between layers that is implicit in feedforward-only networks that use backpropagation for training (Amit, 2019). It has been additionally shown that both local and feedback recurrent connections lead to better performance in more challenging tasks with a better match to neural data, especially during later time points in the response (Lindsay, 2021).

Svanera et al. (2021) showed that a stronger focus on feedback processes can be modelled with a network consisting of an encoder and a decoder structure. FMRI activation patterns during the corner-occlusion paradigm introduced by Smith and Muckli (2010) were more closely related to a CNN with an encoder and decoder than to the feedforward-only VGG16 network. Since the occlusion of one corner leads to a blockage of the feedforward stream in that region of the visual field, feedback signals could be isolated. They point out that some neuronal activity in early visual areas is not related to sensory information but rather to the brains’ inferences about the world transmitted via feedback pathways to early visual areas, and that such top-down effects should be accounted for in biologically constrained neural networks (Svanera et al., 2021).

Our study aims to design a neural network that shows high generalization performance while maintaining biological validity. Traditional CNNs demonstrated difficulties in generalization to novel viewpoints resulting in exponential inefficiencies when dealing with affine transformations (Sabour et al., 2017). Having said that, CNNs were very well able to explain lower-level activation patterns in fMRI studies (Kubilius et al., 2018). On the other hand, CapsNets have a high generalisation ability by using intrinsic spatial relationships to constitute viewpoint-invariant knowledge about an object (Sabour et al., 2017). This was assumed to be a characteristic that is comparable to features of higher-level visual areas of the human brain active during object classification. The original CapsNet (Sabour et al., 2017) is implemented as a shallow network with only three layers which is unlike the human ventral stream which includes computations in a minimum of four layers (Güçlü & van Gerven, 2015; Serre et al., 2005). We, thus, decided to implement a network that incorporates convolution and pooling in lower-level areas as in CNNs, and more complex computations as in CapsNets in higher-level areas. The network comprised V1 and V2 of CORnet-Z (Kubilius et al., 2018) and added the entire CapsNet structure from Sabour et al. (2017) replacing the V4 and IT layers of CORnet-Z. Feedback connections have not been implemented in traditional object recognition ANNs but are implicated in perception and imagery processes. These missing feedback connections might be a major shortcoming that resulted in ANNs deficiency in explaining neurobiological data. That is why feedback was modelled in the proposed network with the help of a decoder network, which closely resembled the encoding layers in reverse order.

The network’s generalization performance for unseen translations was expected to be high and was tested for size, location and rotation generalization. The model was also tested with an occlusion paradigm (Smith & Muckli, 2010) to see whether it functioned within the biological constraint of feedback connections using contextual information. The biological validity was also tested by comparing the network to the imaging data collected by Senden et al. (2019). Feedforward and feedback activations could be distinguished in this data set. Therefore, it was suitable to test the hypothesis that the proposed neural network might model the human ventral stream during perception and imagery.

For a biological comparison to brain imaging data, two approaches were used. Firstly, since capsule activation patterns are assumed to be related to higher-level activation patterns in the human brain during perception and imagery, the fMRI activations in higher-level visual areas were mapped onto the capsule activation patterns of the pre-trained network, and the reconstructions of the trial-wise imagery activations were compared to the input stimuli. In previous work, only reconstructions from perception fMRI activation patterns were analyzed (Qiao et al., 2018). Secondly, correspondences between imagery and perception activations of the CapsNet and the human brain activations were evaluated for lower-level areas. To our knowledge, it was not yet tested whether a network incorporating capsules aligns with activation patterns in the human ventral stream during image perception and imagery. Thus, we tested whether an adapted capsule network which is inspired by biological findings can be utilized to achieve high generalization performance while holding up to relevant biological constraints.

## Methods

### Capsule Networks

#### Structure of the proposed network

The computational model used in this study is a combination of the commonly used CorNet-Z (Kubilius et al., 2018) and the capsule network architecture (Sabour et al., 2017). V1 and V2 were derived from the former. These areas were both represented by a convolutional layer followed by a maxpooling layer. V4 and IT used structures from the capsule network which included another convolutional layer, a primary capsule layer, and the so-called digit capsule layer. The primary capsule layer consists of an additional convolutional layer which utilized the squash function. The digit capsule layer contains four category capsules (4 letters) which consisted each of 16 dimensions. For more details see Figure 1. After this encoding part, a dense layer was added. This dense layer consisted of 64 neurons and used a RELU activation function. The decoder consisted of these calculations in reversed order. For this reason, a function was applied that reverses the squash equation, see Equation 1. Afterwards, a trainable matrix was initialized which served as reversed routing weights.

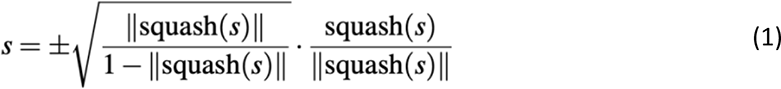

**Figure 1.**
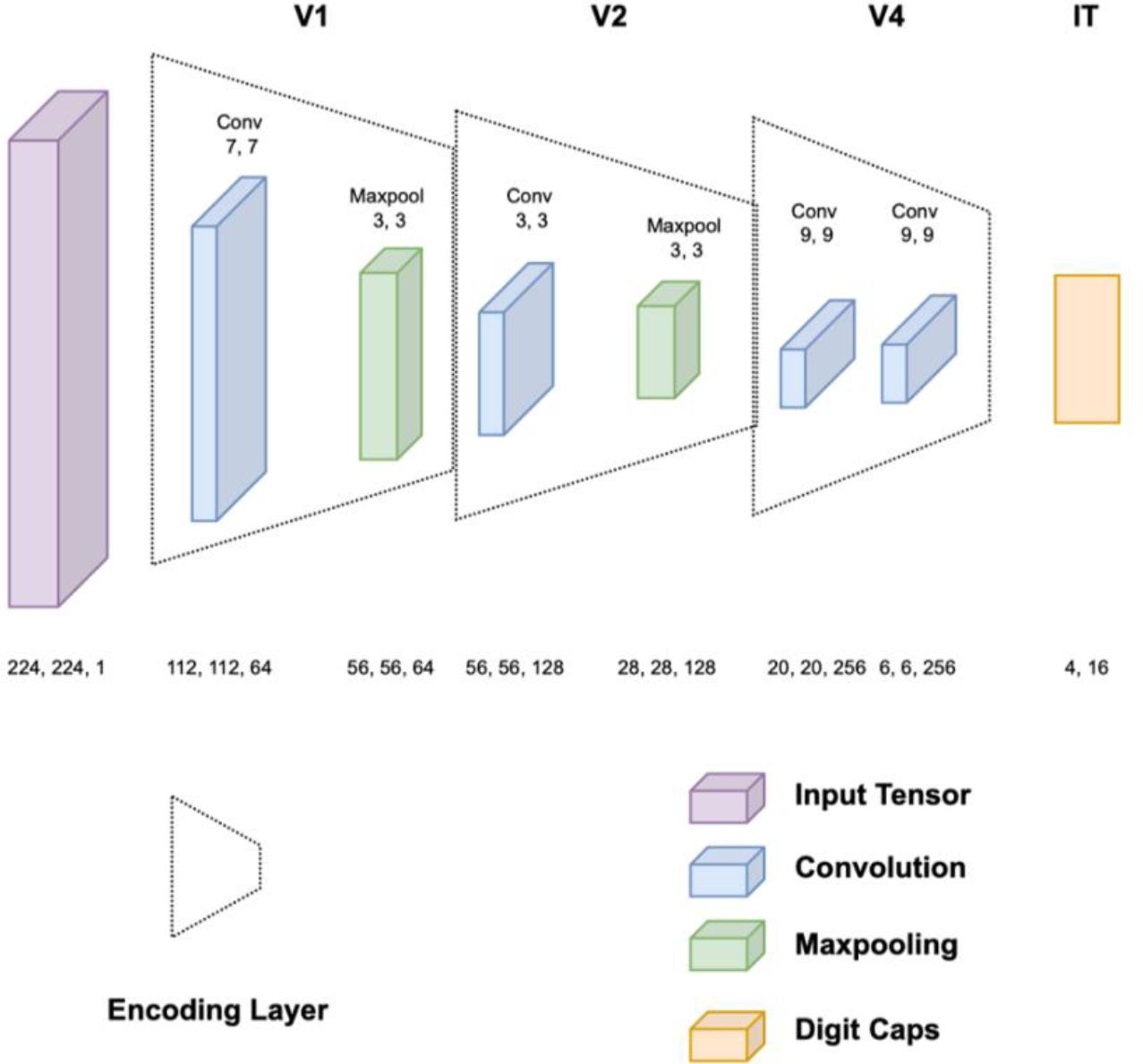
Encoder architecture of the implemented network. V1 and V2 each consisted of a convolutional and a maxpooling layer consecutively. V4 consisted of a convolutional layer and the primary capsule layer which in itself also is a convolutional layer with the squash function applied, whereas IT consists of the so-called digit capsules.

The beforehand calculated unsquashed predictions were then matrix multiplied by the inverse of the reversed routing weights to get to the former predictions of the digit capsule activations. A matrix of the size of the transformation matrix which was needed in the encoder to predict the output of the digit capsules was initialized afterwards. The inverted transformation matrix was multiplied by the predicted activations of the digit capsules. These raw primary capsule activations were then deconvolved with a filter with a size of 9×9 and a stride of two. Another deconvolution layer of size 9×9 and a stride of one was applied. Afterwards, a 2×2 upsampling layer was used. The next layer deconvolved with a filter of size 3×3 and a stride of one. The next upsampling layer had a filter size of 2×2. Lastly, another deconvolution layer with a size of 7×7 and a stride of two was applied.

#### Training and Testing

The network was trained and analysed with the letters C, H, S, and T extracted from the EMNIST dataset (Cohen et al., 2017). Only uppercase letter stimuli were included. These letters were chosen because they are comparable to the stimuli used in the fMRI study by Senden et al. (2019) that was used to test the networks’ biological plausibility. The original CorNet-Z structure expected an input image of 224×224 pixels but the EMNIST stimuli are of size 28×28 pixels which is why they were inserted into matching-sized arrays. To test the generalization performance to different positions and sizes, the letters were rescaled and shifted within a 224×224 pixel space. No other data augmentation was applied.

Altogether, after extraction and balancing of the uppercase letters C, H, S, and T from the data set, 11752 (2938 per category) samples for training, 292 (73 per category) for validation, and 2648 (662 per category) for testing were used. The network used the Adam optimizer. The loss was calculated as the sum of the margin loss and the reconstruction loss. The margin loss was calculated in the same way as in Sabour et al. (2017) (see equation 2).

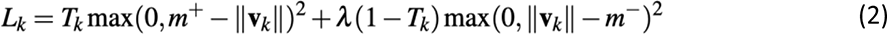

T_k_ was equal to one if the letter of class k is present, otherwise it was zero. As in Sabour et al. (2017), m^+^ was set to 0.9, m^−^ was set to 0.1, and λ was equal to 0.5. The reconstruction loss was calculated as the mean squared difference between the reconstructed picture and the input picture. When adding up the margin loss and the reconstruction loss, the reconstruction loss was scaled down by a factor of 0.0005 to ensure that the margin loss dominated the training. For each test that was run, the network was trained for eight epochs of which the one with the lowest validation loss was saved and used for the analysis. For building this network, Tensorflow 1.7 was used.

#### Location Generalization

The first test of this network entailed resizing the letters to three different possible sizes (50×50, 70×70, 85×85 pixels) and placing them in different positions in a 244×244 pixel image. Training happened on 76.5% of the 14780 possible positions whereas 8.5% were used for validation and the remaining 15% for testing location invariance of the model. The network was trained on the training stimuli in the training positions and tested on separate testing samples in unseen testing positions. To find out about the relevance of the number of dimensions, the number of capsule dimensions was doubled, and the tests were rerun. With the standard amount of 16 dimensions, an additional test was run with half the number of possible locations.

#### Size Generalization

The same network structure was trained in a paradigm that tested whether generalization to unseen sizes of stimuli is possible. Possible letter sizes ranged from 28×28 pixels to 74×74 pixels which accumulates to 46 possible sizes. 76.5% of these sizes were chosen for the training process. 8.5% for validation and the remaining 15% for testing purposes. The classification performance and accuracy can be found in Table 1.

**Table 1.** Generalization Ability of the CapsNet. The CapsNet was trained for different tasks, including size generalization (three different positions with multiple different sizes), location generalization (three different sizes trained and tested in multiple positions with 16 capsules and all sizes), size and location generalization at the same time, as well as rotation generalization (rotation up to 90°in each direction). Training happened on 76.5% of the data with 8.5% for validation and 15% testing data. The respected tested specificities were not presented during training. Classification accuracy and loss calculated as the sum of margin loss and reconstruction loss for each of these tasks are presented. (↑ indicates the more the better, ↓ indicates the less the better).

| Measurement | Size | Location | Size & Location | Rotation |
| --- | --- | --- | --- | --- |
| Classification Accuracy ↑ | 99.62% | 99.81% | 99.51% | 99.77% |
| Loss ↓ | 0.13 | 0.24 | 0.17 | 0.34 |

#### Location and Size Generalization

For this test, the aforementioned different positions, as well as different sizes, were used and matched. Whenever a resized image could not be placed in a certain location, this location was skipped until a fitting location was found (e.g., most stimuli cannot be presented on the edge since they would only be partially presented).

#### Rotation Generalization

With three different locations and three different sizes (50×50, 70×70, 85×85 pixels) of stimuli, a network was trained on rotated images. The images were rotated up to 90°clockwise or counterclockwise in 1° steps.

#### Measures

First and second-order correlations were calculated for each network respectively. First-order correlations are depicted as averages as well as separately by letter. They were calculated by the correlation between the binary letter stimulus and the reconstruction stimulus. The second-level correlation metric was defined by correlating two vectors, one depicting the pairwise correlation between physical letter stimuli and the other depicting pairwise correlations between reconstructions. Additional measures were used to assess the similarity between the images, namely root mean squared error (RMSE), peak signal-to-noise ratio (PSNR), structural similarity index measure (SSIM), and euclidean distance.

#### Occlusion Paradigm

To test whether the model activations also relate to the results from Smith and Muckli (2010), an adapted occlusion paradigm was used. Smith and Muckli (2010) showed that fMRI activation relating to non-stimulated regions of the visual field could be used to correctly classify the presented category above chance with a support vector machine (SVM) classifier that was trained on activation patterns elicited during perception.

To test whether this holds for the suggested CapsNet, 70×70 pixel stimuli from the test data set were presented in the centre of the image. One of the corners was occluded which resulted in an occlusion area of 35×35 pixels. The reconstructions of the complete stimuli were cut out in the relevant area, an edge of five pixels was left towards the sides that would be later-on non-occluded. The SVM classifier was trained on these corner activations. Afterwards, the occluded corner reconstructions were used to predict their respective stimuli categories with the pre-trained classifier.

### Functional Magnet Resonance Imaging Data

Six participants (two female, four male; average age of 30.7 years) underwent ultra-high field (7 Tesla) fMRI. For more details on the data set and the processing steps, see Senden et al. (2019).

#### Stimuli and Task

Three training sessions were completed by each participant prior to a single scanning session which consisted of four experimental (imagery) runs, one control (perception) run and a pRF mapping run. Training sessions lasted about 45 min and took place one week before scanning. Each training session consisted of the participant seeing one of four white letters (C, H, S, or T) enclosed in a white square guide box on a grey background and a red fixation dot in the centre of the screen. With the onset of visual presentation, an auditory stimulus was presented that comprised three low tones and one high tone. Specific tone patterns were associated with a visually presented letter randomly assigned per participant. At the beginning of each training session, each of the four letters was presented for 3000 ms together with their respective tone pattern. During one training run, each participant completed 16 pseudo-randomly presented trials. Each training session consisted of two training runs in which the reference letters with their tone patterns were presented upon beginning, and two training runs without reference letter presentation. At the beginning of each trial, after presenting the letter and the tone for 3000 ms, the letter faded out, having disappeared after 5000 ms. Afterwards, the fixation dot changed colour to orange and participants had to maintain a vivid mental image of the presented letter. After 18 s of imagery, the fixation dot turned white and three white probing dots appeared within the guide box, initiating probing. The dots were either located within the letter shape or outside of it and participants had to indicate which one it was. The fixation dot turned green if the answer was correct or red for incorrect answers.

Imagery runs differed from the training runs in probing phase and timing of trial phase. Referencing still took place but was followed by trials that did not contain visual stimulation besides the fixation dot and the guide box. Imagery phases were initiated by tone pattern presentation and the fixation dot turning orange. Imagery phases lasted for 6 s. Each experimental run contained 32 normal trials, with two additional catch trials with probing. No visual feedback for the probing phase was given.

The perception run was used to measure the brain activation patterns in visual areas during perception of the letters. The same timing parameters as in the experimental run were used. No reference or probing phases were needed. Letters were presented for the duration of 6 s while their shape was filled with a flickering checkerboard pattern. No tone patterns were played during the perception run. For pRF mapping, a bar aperture revealing a flickering checkerboard pattern was presented in four orientations. In twelve discrete steps, the bar covered the entire screen for each orientation. Within each orientation, the sequence of steps was randomized, and each orientation was presented six times.

#### Processing of fMRI Data

To test the biological plausibility of the network, the voxel activation patterns of early visual areas V1 and V2 with ROIs from Senden et al. (2019) were used. Furthermore, activations that were extracted from early visual areas, pRF mapped and run through a pre-trained autoencoder were utilized for the analysis. These reconstructions had a size of 150×150 pixels. The relevant part of the reconstructed stimuli from the CapsNet was cut and rescaled in differently sized patches due to the different sizes of the layers. Additionally, activation patterns from the higher visual areas localized by significant activation patterns averaged over all letters were extracted.

#### Translating fMRI Data into Capsule Space

Since the capsule network would have at its highest layer 4×16 = 64 dimensions and higher visual areas indicated 734 relevant voxels, the voxels had to be translated into the digit capsule space. For that purpose, a simple three-layered deep neural network was used. Inspired by Qiao et al. (2018), the first two layers were dense layers with the RELU activation function, whereas the last layer used the squash function adapted from the CapsNet. The first layer contained 256, the second layer 128, and the last layer 64 units—these 64 units were eventually compared to the highest layer of the CapsNet. The model was trained on the perception data, with seven trials from each letter serving as training data, and one trial used as validation data. This network was trained with the Adam optimizer and ran for ten epochs. This pre-trained network was then used to predict the capsule activation patterns for each imagery trial separately. No additional training took place.

#### Representational Similarity Analysis

To compare the activation patterns of the neural network and the fMRI activation patterns, a representational similarity analysis (RSA) was used (Kriegeskorte, Mur, & Bandettini, 2008). This method uses second-level multidimensional scaling to relate the representations of information in different modalities to each other. Representational dissimilarity matrices (RDMs) were calculated with the help of the Python package rsatoolbox. They were calculated for V1 and V2 encoder and decoder activations when the artificial letter stimuli from the fMRI study served as input. RDMs based on V1 and V2 decoding activations of the CapsNet were also calculated for reconstructions from remapped activation patterns from higher-level visual areas. Dissimilarities were measured with 1−correlation. These RDM patterns were then used for the RSA. The RSA compared with the help of correlations the activation patterns in RDMs of the CapsNet with the fMRI data, during perception and imagery.

## Results

### Location Generalization

The results of a network that was trained to classify images in different locations are shown in Tables 1, 2, 3, and Figure 2a. The second-order correlation of one hundred stimuli amounted to .88. Reconstructions of ten example stimuli are depicted in Figure 3. When the number of dimensions was doubled to 32, the classification performance reached 99.81% with a loss of 0.21. When the amount of possible locations was halved to 7390 and the number of dimensions per capsule was kept at 16, the classification accuracy amounted to 99.73% and the loss reached 0.24.

**Figure 2.**
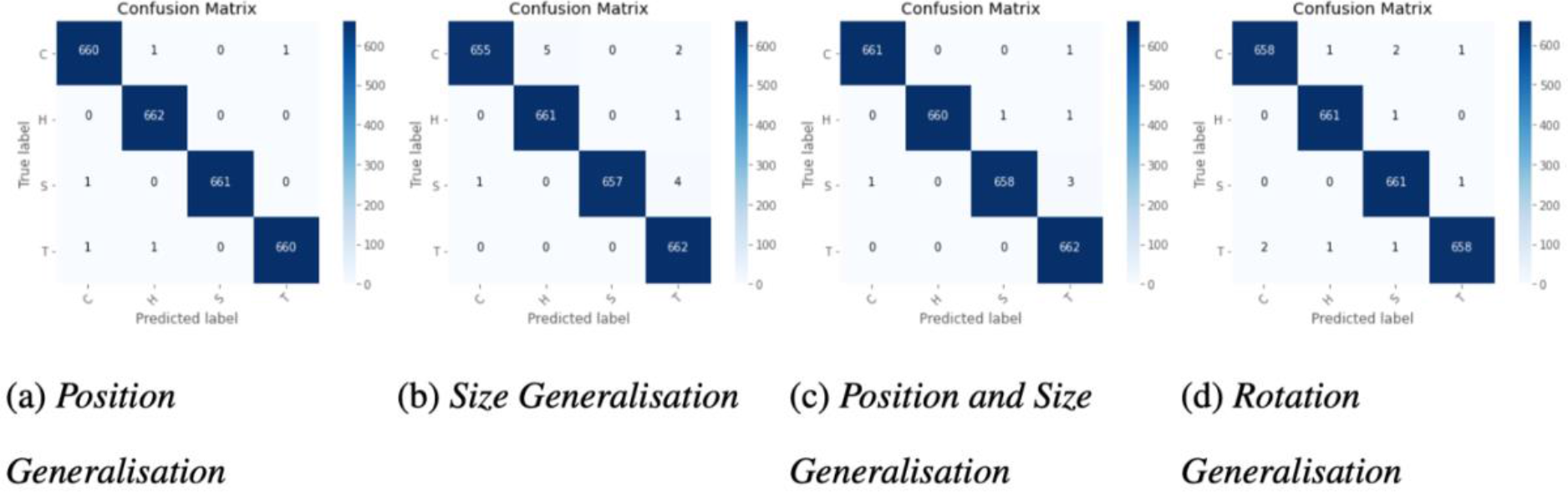
Classification Performances for Generalization Tasks. Classification performances for the differently trained networks (position generalization with three sizes in multiple positions, size generalization with three positions in multiple sizes, a combination of both, and rotation generalization with rotations up to 90°in either direction) depicted in a confusion matrix with absolute numbers of correct classifications. True labels are depicted on the y-axis and predicted labels are depicted on the x-axis. Classification performances were high in every condition.

**Table 2.**
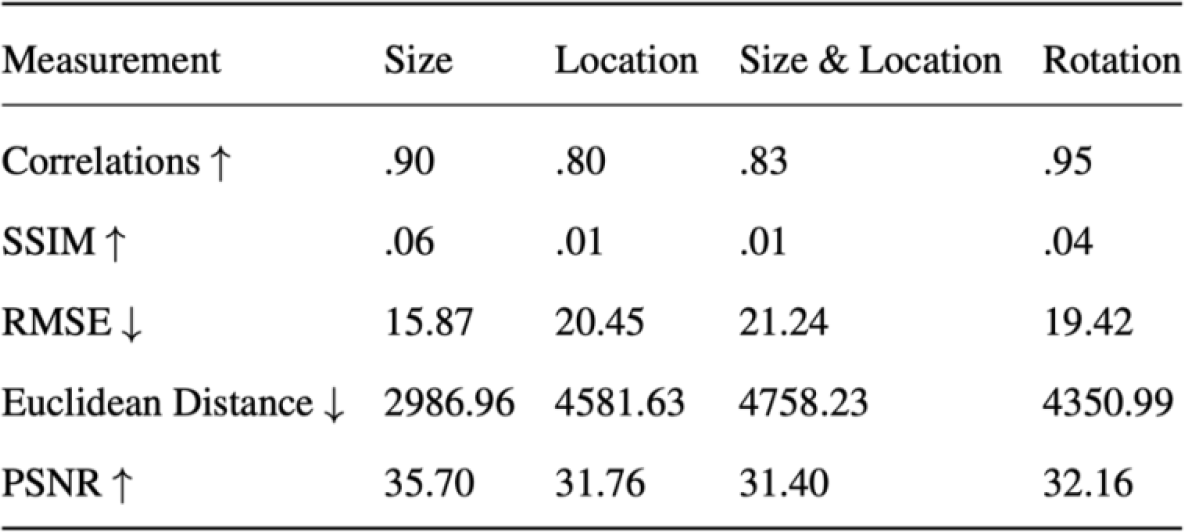
Similarity Measures between Input and Reconstructions. First-level reconstruction quality measures indicating similarity between input and reconstruction images averaged for all letters. Results are represented for each training process for different forms of generalization individually (SSIM = Structural Similarity Index, RMSE = Root Mean Square Error, PSNR = Peak Signal-to-Noise Ratio, ↑ indicates the more the better, ↓ indicates the less the better).

**Figure 3.**
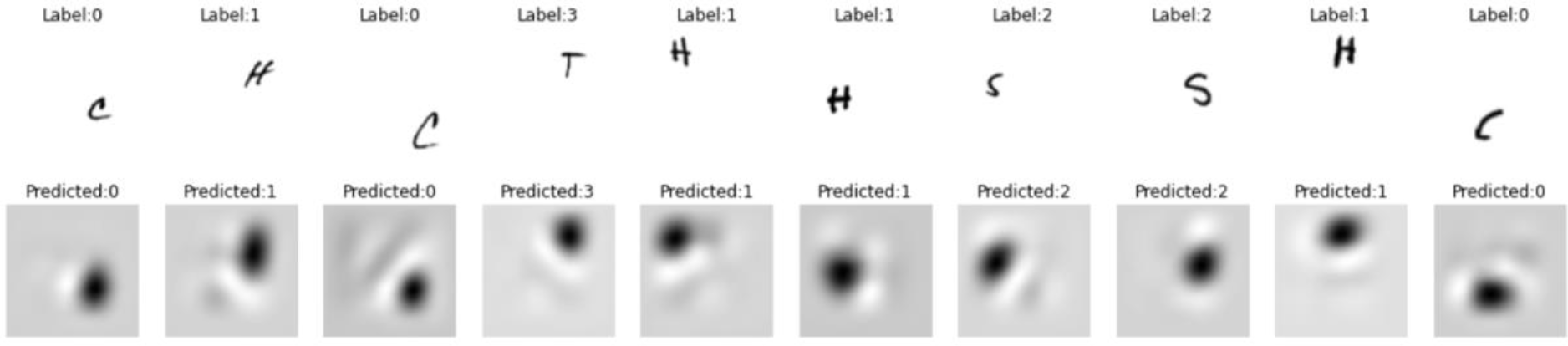
Reconstructions for Location Generalization. Input and reconstruction of CapsNet trained on different positions of letter stimuli. Testing and training positions were entirely independent. Reconstruction letters are not distinguishable. The labels indicate which category the stimulus belonged to as well as the predicted label from the network classification (0 = C, 1 = H, 2 = S, 3 = T).

**Table 3.** Similarity Measures for Location Generalization Task. First-order measures between reconstructed letters and physical stimulus per letter for the network trained on location generalization (3 different sizes in multiple locations in the picture, trained and tested locations were entirely independent). (SSIM = Structural Similarity Index, RMSE = Root Mean Square Error, PSNR = Peak Signal-to-Noise Ratio, ↑ indicates the higher the more similar, ↓ indicates the lower the more similar).

| Measurement | C | H | S | T |
| --- | --- | --- | --- | --- |
| Correlations ↑ | .78 | .83 | .79 | .77 |
| SSIM ↑ | .02 | .01 | .01 | .01 |
| RMSE ↓ | 21.57 | 22.10 | 21.99 | 19.78 |
| Euclidean Distance ↓ | 4831.83 | 4949.60 | 4926.03 | 4431.07 |
| PSNR ↑ | 31.31 | 31.11 | 31.15 | 32.08 |

### Size Generalization

The results testing whether the network can classify images in unseen locations can be found in Table 4. For reconstruction results see Figure 4. The classification performance for stimuli with unseen sizes is depicted in Figure 2b. Second-order correlation for 100 stimuli averaged up to 0.76. When the stimuli from the fMRI study were used as input, the classification worked perfectly, reconstruction results can be seen in Figure 5.

**Figure 4.**
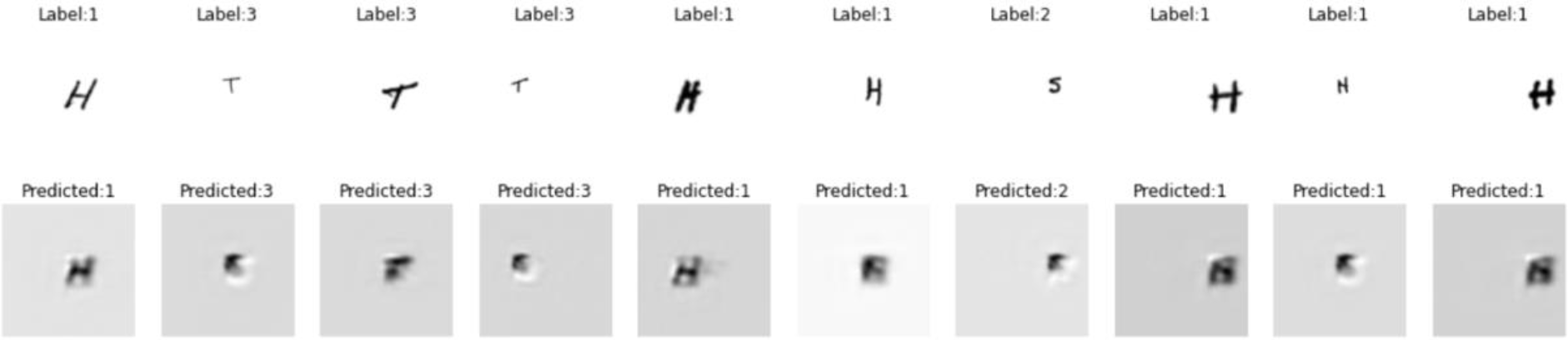
Reconstructions for Size Generalization. Input and reconstruction of CapsNet trained on different sizes of letter stimuli. The depicted sizes were only used as part of the test data set. The labels indicate which category the stimulus belonged to, and the predicted labels indicate which classification the network allocated to the letter (0 = C, 1 = H, 2 = S, 3 = T).

**Table 4.** Similarity Measures for Size Generalization Task. First-order measures between reconstructed letters and physical stimuli per letter for the network trained on size generalization. Different sizes were presented in three different positions. None of the sizes used in the test data set were part of the training data set. (SSIM = Structural Similarity Index, RMSE = Root Mean Square Error, PSNR = Peak Signal-to-Noise Ratio, ↑ indicates the higher the more similar, ↓ indicates the lower the more similar).

| Measurement | C | H | S | T |
| --- | --- | --- | --- | --- |
| Correlations ↑ | .91 | .93 | .91 | .91 |
| SSIM ↑ | .08 | .09 | .07 | .09 |
| RMSE ↓ | 15.66 | 16.56 | 15.89 | 14.19 |
| Euclidean Distance ↓ | 3508.20 | 3710.44 | 3559.46 | 3178.94 |
| PSNR ↑ | 34.18 | 33.66 | 34.04 | 35.03 |

**Figure 5.**
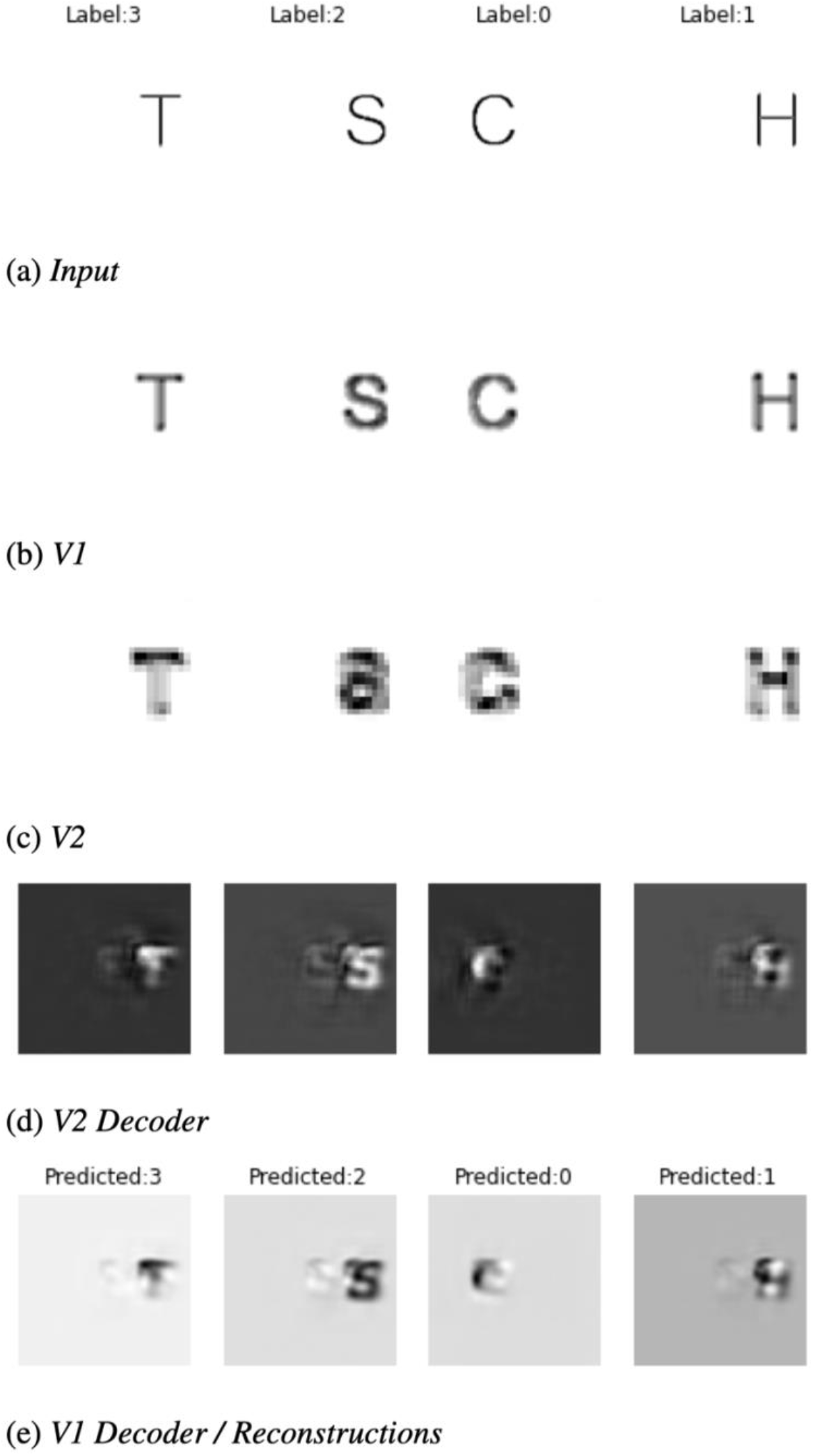
Reconstructions of Artificial Letter Stimuli. Activations in different layers of the network when the CapsNet was trained on different sizes with three different positions and tested on the artificial stimuli that were used as part of the fMRI study that these activations were compared to. Depicted are the input to the pre-trained CapsNet, V1, V2, V2 decoder, and the V1 decoder activations. The V1 decoder activation is also seen as the reconstruction activation. The labels indicate which category the stimulus belonged to, and the predicted labels indicate which classification the network allocated to the letter. All four letter stimuli were correctly classified (0 = C, 1 = H, 2 = S, 3 = T).

### Location and Size Generalization

The network was tested on a paradigm to see whether size and position generalization can be trained at the same time. Results can be found in Tables 1, 2, and 5 as well as Figure 2c. Reconstructions are seen in Figure 6. The second-order correlation of one hundred stimuli averaged up to .71.

**Table 5.** Similarity Measures for Location and Size Generalization Task. First-order measures between reconstructed letters and physical stimuli per letter for the network trained on location and size generalization. Different sizes and locations were used for training, not restricted to an amount of three on any dimension. The depicted results stem from testing positions and sizes that were not included in the training data set. (SSIM: Structural Similarity index measure, RMSE: Root Mean Squared Error, PSNR: Peak Signal-to-Noise-Ratio, ↑ indicates the higher the more similar, ↓ indicates the lower the more similar).

| Measurement | C | H | S | T |
| --- | --- | --- | --- | --- |
| Correlations ↑ | .67 | .70 | .71 | .58 |
| SSIM ↑ | .01 | .01 | .01 | .01 |
| RMSE ↓ | 17.60 | 17.78 | 17.60 | 16.10 |
| Euclidian Distance ↓ | 3942.86 | 3983.83 | 3941.96 | 3606.54 |
| PSNR ↑ | 33.33 | 33.19 | 33.28 | 34.14 |

**Figure 6.**
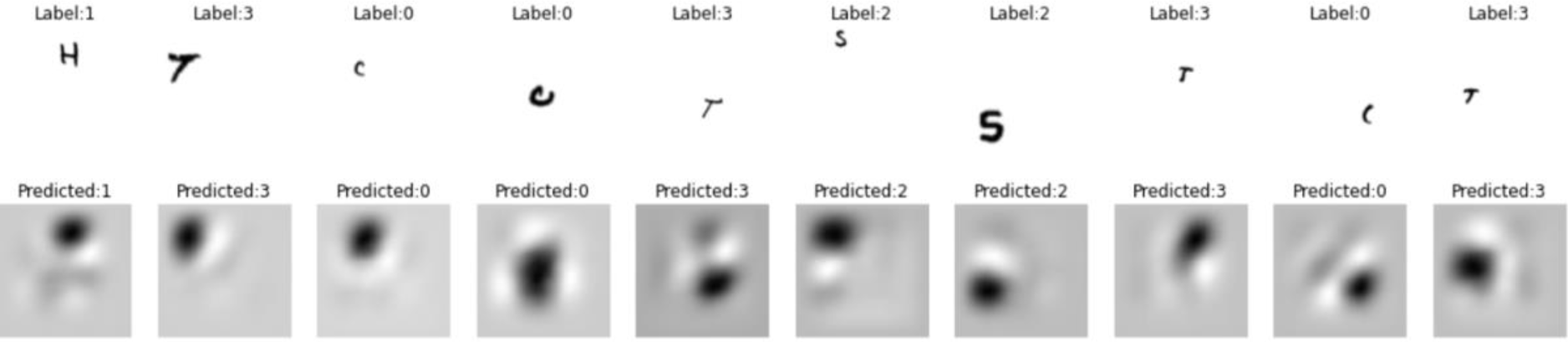
Reconstructions from Size and Location Generalization. Input and reconstruction of CapsNet trained on different positions and different sizes of letter stimuli. The labels indicate which category the stimulus belonged to, and the predicted labels indicate which classification the network allocated to the letter (0 = C, 1 = H, 2 = S, 3 = T).

### Rotation Generalization

Classification accuracy and loss for the rotation generalization task can be found in Table 1. First-order measures of similarity are depicted in Table 2. The second-order correlation amounted to .75 over a sample of 100 stimuli. The reconstructions are shown in Figure 7.

**Figure 7.**
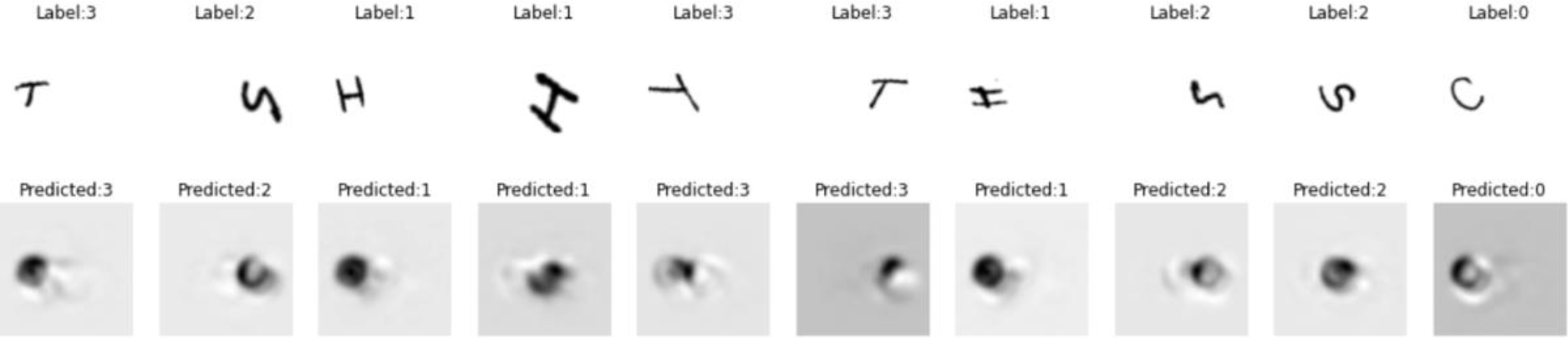
Reconstructions from Rotation Generalization. Input to CapsNet trained on different rotations with three different sizes and positions, depicting input, and reconstruction. The rotations ranged from -90°to 90°. The labels indicate which category the stimulus belonged to, and the predicted labels indicate which classification the network allocated to the letter (0 = C, 1 = H, 2 = S, 3 = T).

**Table 6.** Similarity Measures for Rotation Generalization Task. First-order measures between reconstructed letters and physical stimuli per letter for the network trained on rotation generalization with three90°cdifferentpositionsandsizes.The rotations varied from clockwise to counterclockwise. The testing rotations were not part of the training rotations data set. (SSIM: Structural Similarity index measure, RMSE: Root Mean Squared Error, PSNR: Peak Signal-to-Noise-Ratio, ↑ indicates the higher the more similar, ↓ indicates the lower the more similar).

| Measurement | C | H | S | T |
| --- | --- | --- | --- | --- |
| Correlations ↑ | .93 | .92 | .92 | .90 |
| SSIM ↑ | .04 | .04 | .04 | .03 |
| RMSE ↓ | 24.09 | 27.41 | 25.13 | 24.07 |
| Euclidian Distance ↓ | 5395.86 | 6140.17 | 5630.0 | 5392.04 |
| PSNR ↑ | 30.38 | 29.22 | 29.98 | 30.36 |

### Testing with Occlusion Paradigm

Using a model that was trained on three different locations with multiple sizes, it was tested whether similarly to fMRI activations in Smith and Muckli (2010), non-stimulated regions of the network elicited activation patterns that made it possible that an SVM classifier evaluated the presented category above chance level. Some examples of reconstructed stimuli from complete and occluded stimuli can be seen in Figure 8. Stimuli which had an occluded corner were fed into the network and the reconstructions were extracted. When this reconstruction was then again fed into the network for classification purposes, the accuracy for the classification was still high above chance. The reconstructions from samples that were occluded in the lower left corner were correctly classified in 60.16% of cases, whereas when the lower right corner was occluded, only 43.24% were correctly classified any longer indicating that the lower right corner contained more relevant information than the upper left one. The upper left occlusion led to an accuracy of 32.68%, and the upper right corner occlusion of 42.03%. The confusion matrices can be found in Figure 9. The SVM classifier that was trained on the reconstructed corner area from whole stimuli, was then used to predict the stimulus category from the reconstruction of occluded pictures from the other half of the testing data set. Results for correct classification by the SVM classifier can be seen in Figure 10. Even though the corner did not contain any information in the input stimulus for occluded stimuli, the reconstruction showed a classifiable pattern of activation. 96.79% were correctly classified. Training on perceptual stimuli and testing on reconstructed stimuli from occluded letters did not lead to meaningful results.

**Figure 8.**
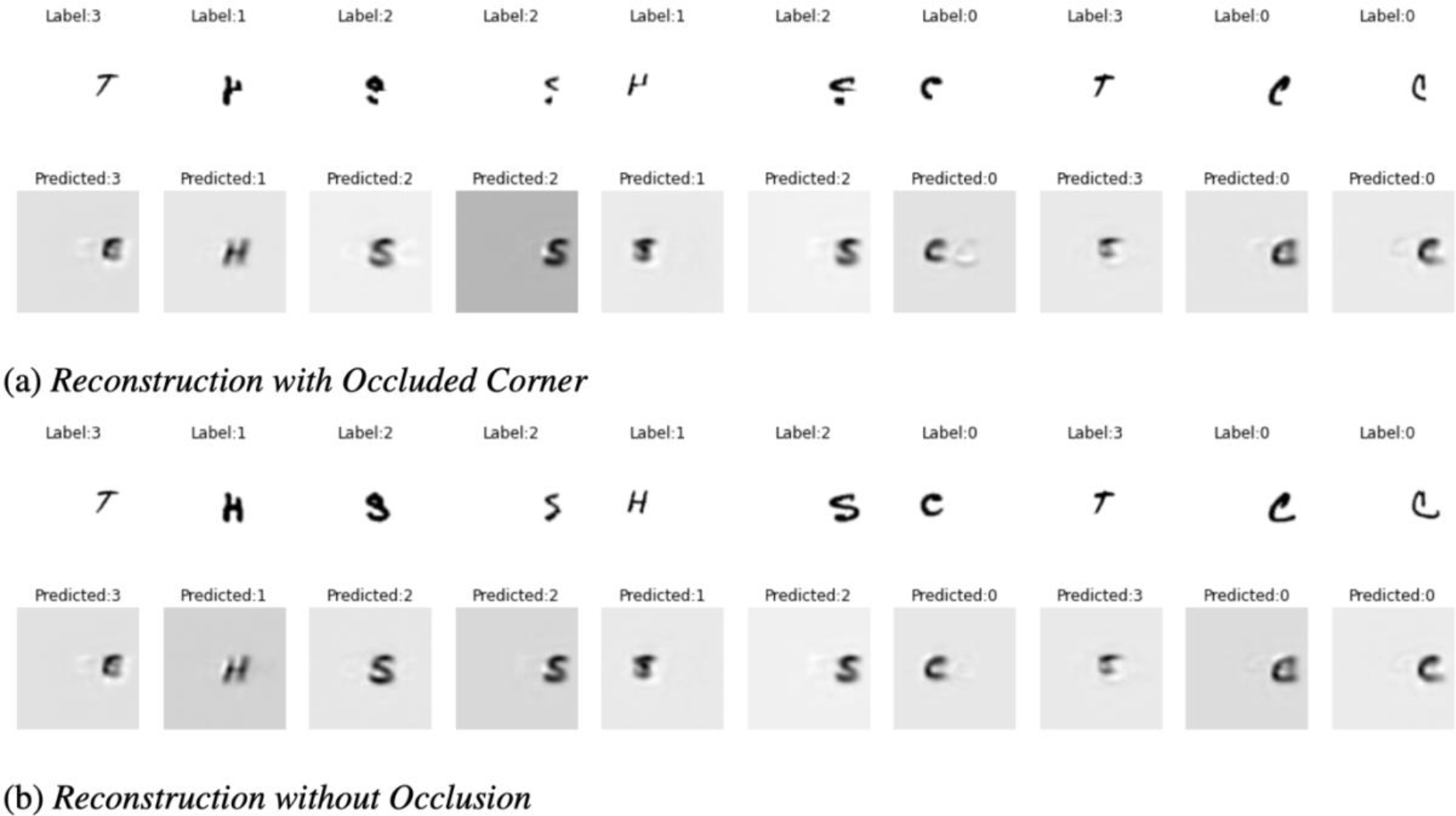
Reconstructions for Occlusion Paradigm. Pictures that occluded the lower right corner were inserted in the network which was trained on three different positions with different sizes. To make the analysis more concise, only 70×70 pixel letter stimuli were used here. Reconstructions from the occluded pictures can be compared to the reconstructions from non-occluded pictures. Labels indicate the category to which the stimulus belongs to, whereas the predicted label indicates how the network classifies the stimulus (0 = C, 1 = H, 2 = S, 3 = T).

**Figure 9.**
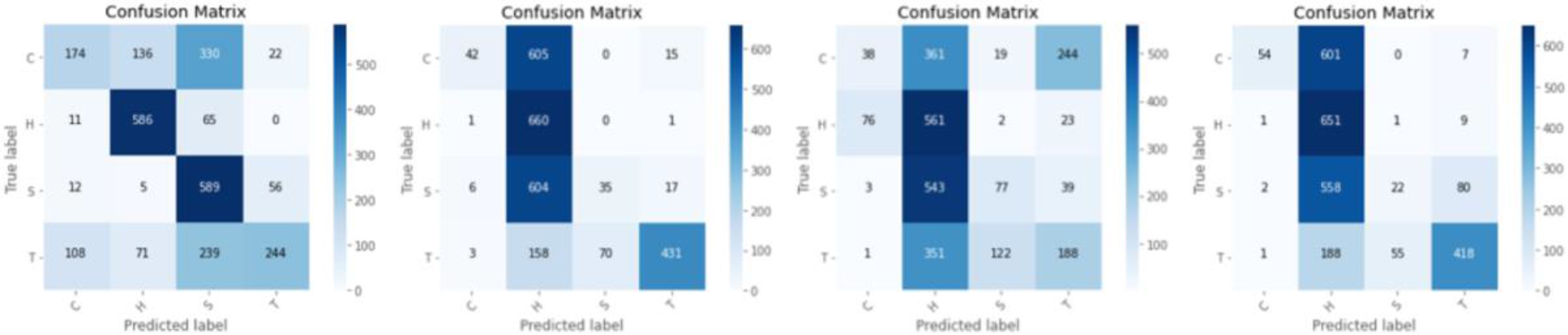
Classification Performance for Reconstructions from Occluded Stimuli. The four different corners of the letter images were occluded. The reconstructed stimuli were fed through the same network again and classifications are depicted in a confusion matrix. Strong bias towards the ’H’ stimulus could be seen. Classification for each of the occluded corners is above chance level. On the x-axis, the predicted label is shown whereas the y-axis shows the true label.

**Figure 10.**
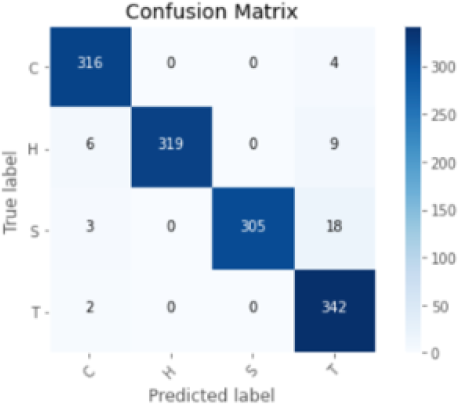
Classification for Corners Reconstructed from Occluded Stimuli. This picture shows the SVM classification of the reconstruction of occluded stimuli. The stimuli were occluded in the lower right corner. This classifier was formerly trained on the reconstructed corner of the complete stimuli. Half of the testing stimuli were used for training and the other half for testing. This high classification performance indicated that during reconstruction even in areas which relate to input areas where no information was shown, relevant activation could be seen.

### Relating the CapsNet with the fMRI Data

#### Reconstructing Higher-Level Imagery Activations

When the fMRI data was mapped with the help of a neural network onto the capsule space, a validation loss of 0.03 was reached. The validation trial reconstructions are depicted in Figure 11.

**Figure 11.**
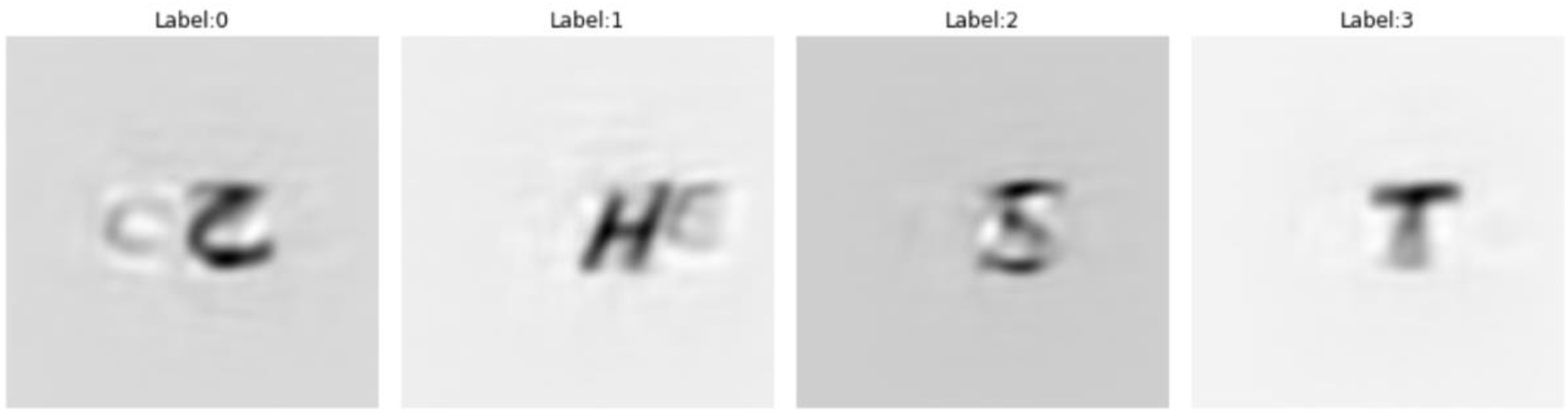
Perception Trial Reconstructions. Reconstruction results when perception voxel activations of the validation trials were mapped onto capsule activation patterns with the help of a three-layered deep neural network and then inserted into the CapsNet. This pre-trained three-layered network was then used to map the imagery higher-level activation patterns onto the capsule space.

It was seen that correlations and SSIM of the entire pictures did not yield any relation between the reconstructed stimuli and the presented stimuli. RMSE and Euclidean distance showed the lowest and therefore best-matching scores for three out of four stimuli. Only the letter S was not closest related to the actual S stimulus for each of these indices. When taking the first dimension of each category of each remapped stimulus to see whether the network learned the different classes correctly, it became clear that the highest value for the presence of a specific category stimulus was not of importance for the network. This value only reached chance level. On the other hand, when correlating the capsule activations with the expected capsule activations for the imagery stimuli, 38 out of 128 stimuli were strongest correlated with their expected capsule activations compared to the other category capsule activations (binomial test, p = .09). Additionally, the predicted capsule activations for each trial were fed through the decoder of the pre-trained CapsNet. Some examples of reconstruction results can be seen in Figure 12. The reconstructions from single trials were significantly the highest correlated with the respective stimulus (binomial test, 128 trials, 47 reconstructions with highest correlation with respective letter stimulus, 36.72%, p < .001). When using just a cutout part (100×120 pixels) of the letter stimuli, 51 (39.84%) of the stimuli were higher correlated with their respective letter stimulus compared to their correlation with the other physical stimuli. An overview of these strongest correlations for the full images and the cutout image can be found in Figure 13. When using Euclidean distance as a measurement, 67 reconstructed stimuli (50.78%) showed the smallest distance to the respective physical letter stimulus for the entire images and 62 (46.88%) did so for the smaller cutout area. Results for SSIM were highly similar to the correlational results. The results for RMSE as well as PSNR were comparable to the Euclidean distance plots. Relevantly, these results were highly susceptible to the respective pixel area that was chosen; thus, these results should be interpreted with caution.

**Figure 12.**
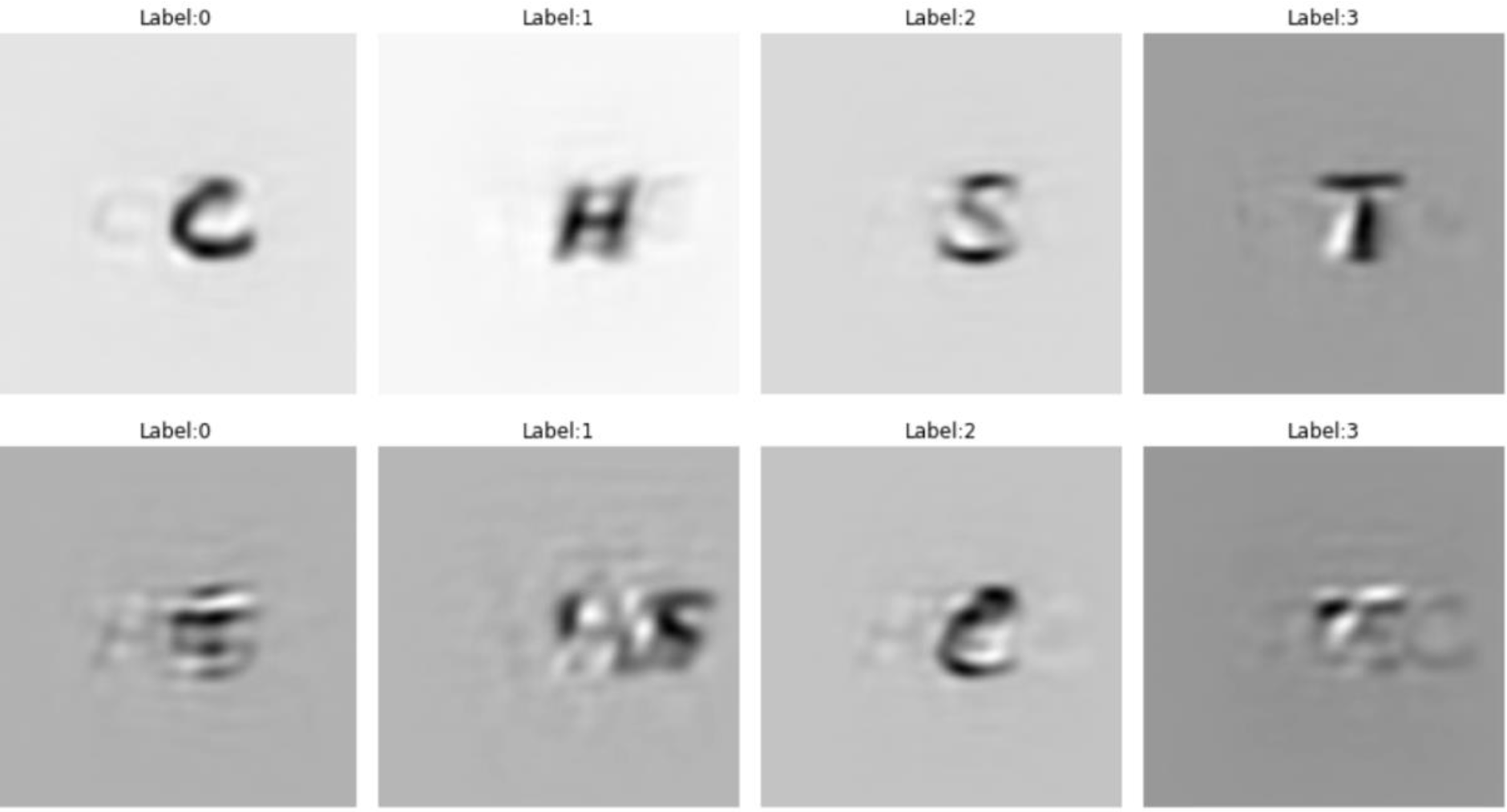
Exemplary Reconstructions from Imagery Trials. Exemplary reconstruction results when imagery voxel activations were mapped onto capsule activation patterns with the help of a three-layered deep neural network and then inserted into the CapsNet. The first row depicts subjectively identifiable letters whereas the second row depicts examples of reconstructions with restricted identifiability. The labels indicate the respective stimulus group (0 = C, 1 = H, 2 = S, 3 = T).

**Figure 13.**
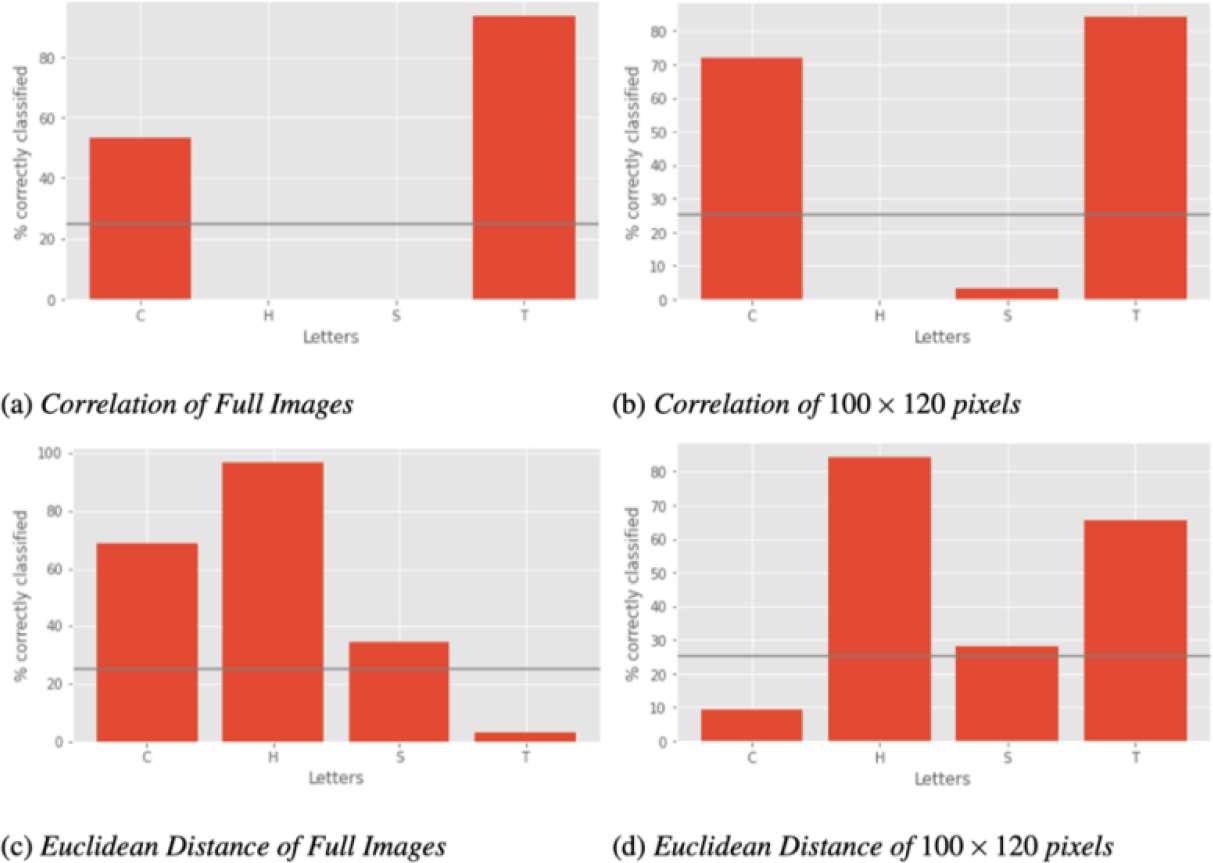
Imagery Reconstruction Evaluation. Evaluation of reconstructions from higher-level imagery activations. The number of strongest correlations, and lowest Euclidean distances of reconstructed letters with respective letter stimulus separated by letter. Depictions for full images and image cutouts of 100×120 pixels. The measures yielded different results. The grey line represents a chance level of classification at 25%.

As a final evaluation, the reconstructed pictures were fed through the network to see whether the network itself would classify the pictures according to their respective expected categories. The network classified 60.94% (70 correctly classified stimuli) of the stimuli according to their imagined stimulus. The confusion matrix for the correct and incorrect classifications can be found in Figure 14.

**Figure 14.**
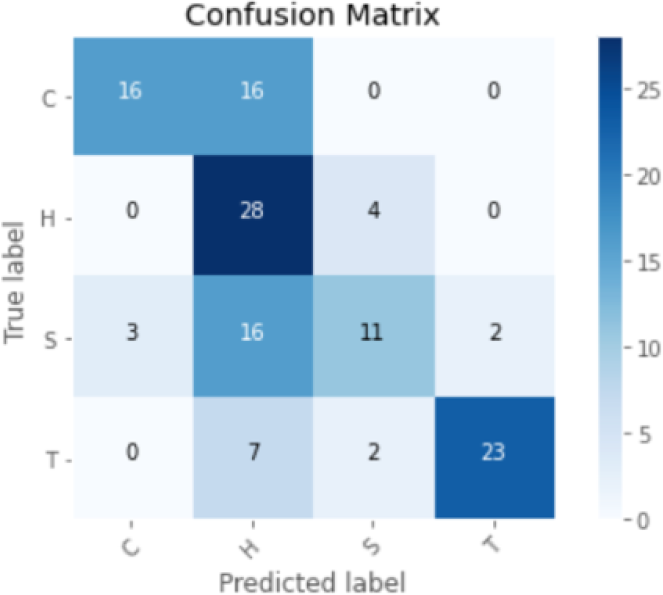
Classification Performance for Reconstructed Imagery Stimuli. Classification accuracy for reconstructed imagery stimuli. The CapsNet which was trained on three positions and multiple different sizes was used for this analysis. The x-axis depicts the predicted labels and the y-axis the true labels. The correct classification reached above chance level which indicates that it is possible to reconstruct images from higher-level visual area activations. 60.94% of reconstructions were correctly classified.

#### Comparing Lower-Level Activations of the CapsNet to fMRI Activations

To calculate RSA similarity measures, RDMs for V1 and V2 from the CapsNet encoder and decoder, as well as V1 and V2 raw voxel activations of the fMRI activations during perception were calculated (see Figure 15). The same calculations were done for the fMRI imagery activation patterns for V1 and V2 as well as the reconstructed activation patterns from remapped voxel-to-capsule activations (see Figure 16).

**Figure 15.**
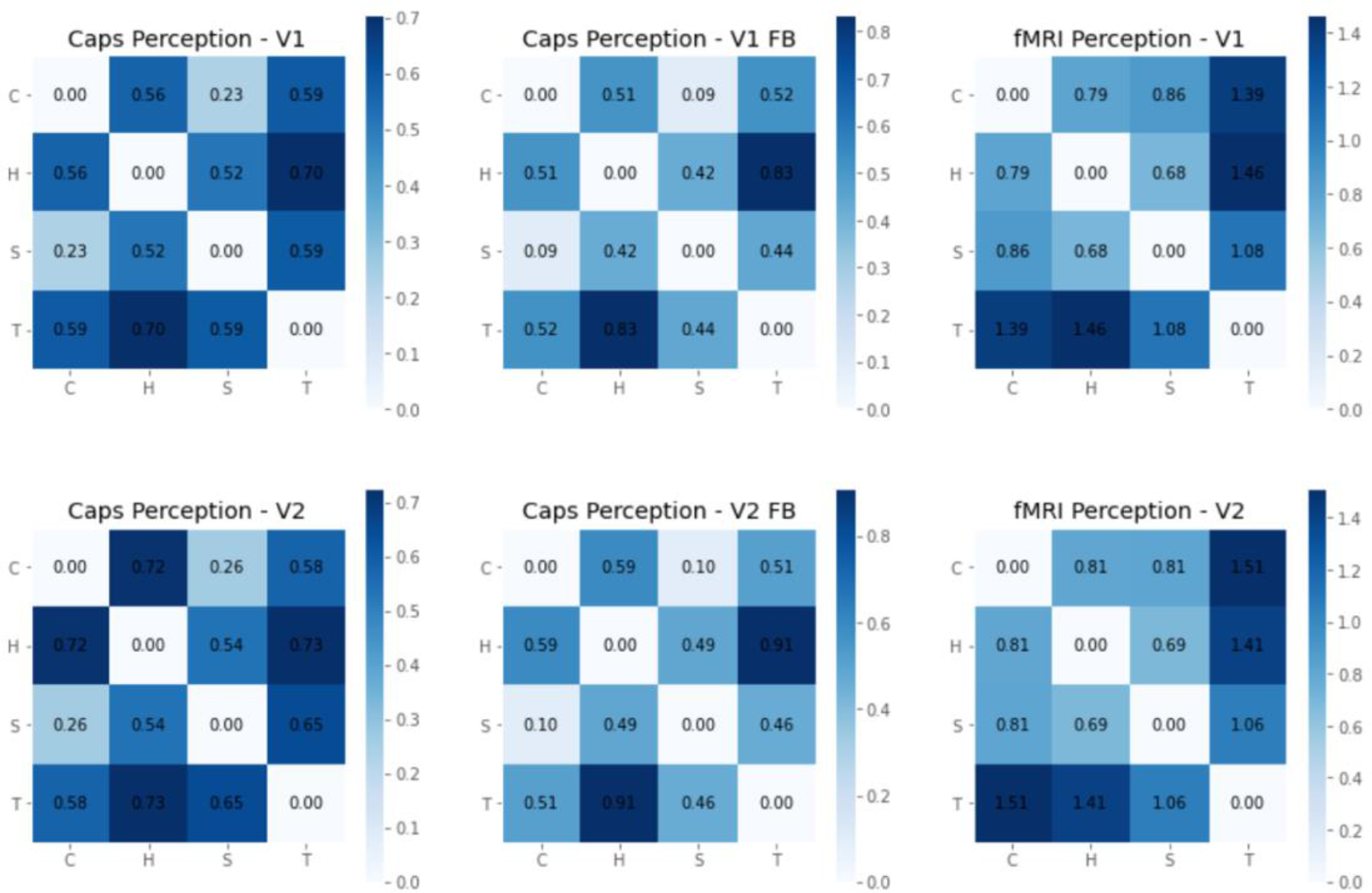
Representational Dissimilarity Matrices from Perception Activation Patterns. These Representational Dissimilarity Matrices were calculated for CapsNet activation patterns relating to V1 and V2, as well as the decoding layers V1 and V2. Additionally, the same measure was calculated for the fMRI activations. The dissimilarities are calculated by 1-correlation between two conditions. Higher values indicate stronger dissimilarities between conditions.

**Figure 16.**
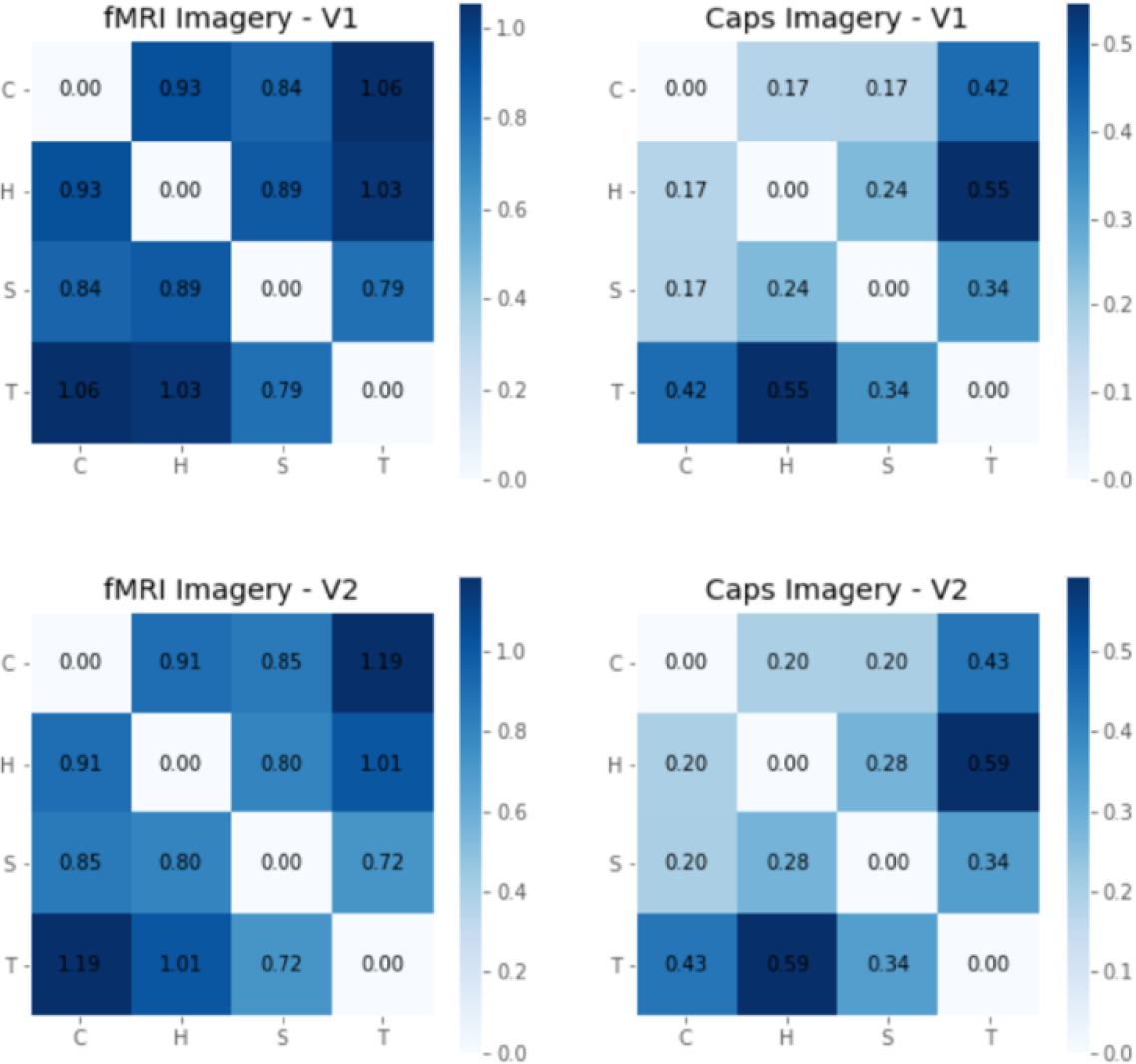
Representational Dissimilarity Matrices from Imagery Activation Patterns. The Representational Dissimilarity Matrices were calculated for CapsNet decoder layers V1 and V2 for reconstructions from imagery activations and fMRI raw voxel activations of layers V1 and V2 during imagery. The dissimilarities are calculated by 1-correlation between two conditions. Higher values indicate stronger dissimilarities between conditions.

To compare these lower-level activation RDMs, Kriegeskorte et al. (2008) suggested RSA analysis. This was applied to compare the activation patterns between the modalities (fig. 17). The correlations for all RDMs were high, not differentiating between representations for V1 and V2. It would have been expected that the fMRI V1 activations related higher to CapsNet V1 representations than to CapsNet V2 representations. The encoder was expected to be more strongly related to the perception fMRI activations and the decoder to the imagery activations. More stimuli categories would be needed to draw conclusions about the representation patterns of the CapsNet and fMRI activation patterns.

**Figure 17.**
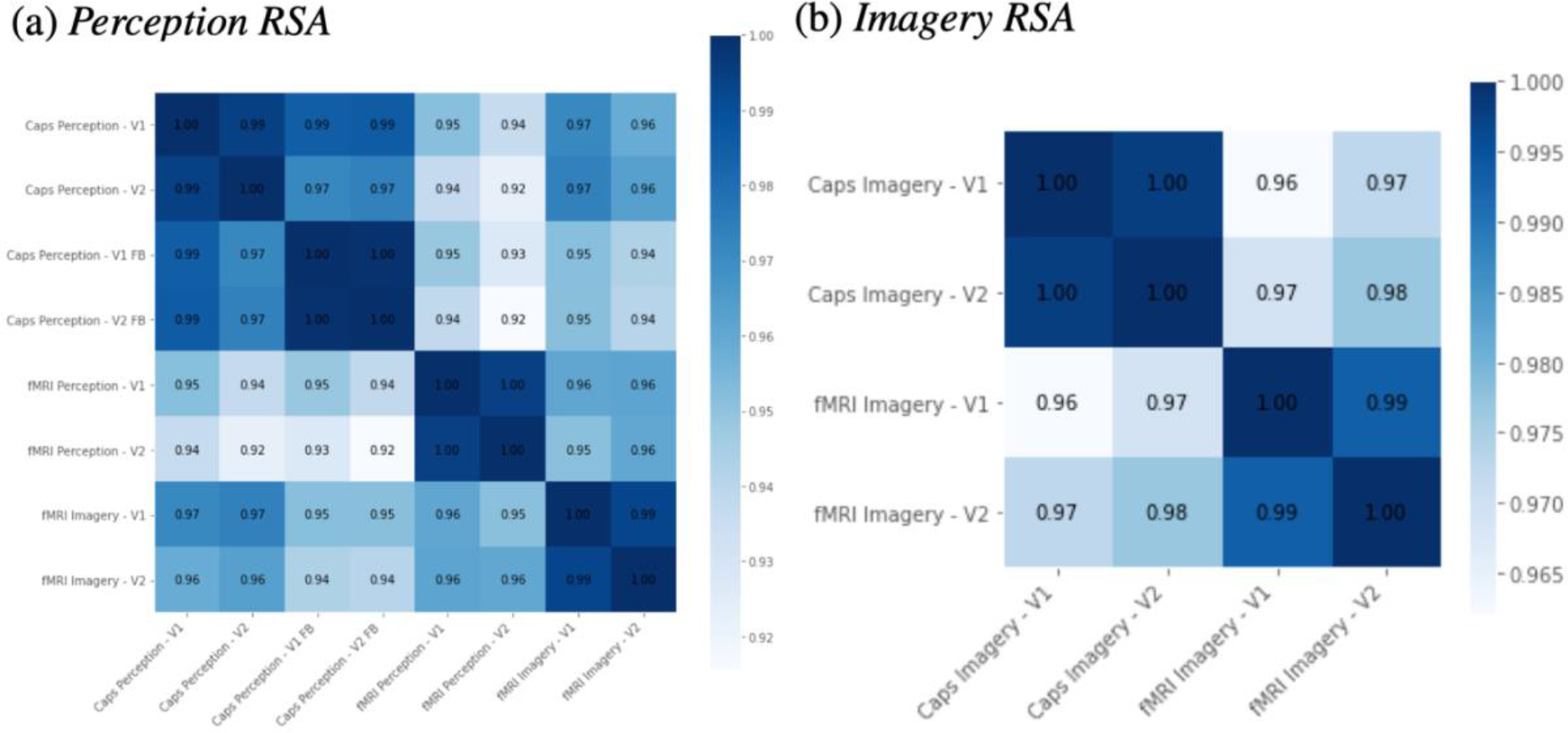
Representational Similarity Analysis of Dissimilarity Matrices. The Representational Similarity Analysis is based on correlations between Representational Dissimilarity Matrices (RDM). This analysis compares RDMs calculated for fMRI raw voxel activations of layers V1, V2, and RDMs based on CapsNet layer V1 and V2 activation patterns, for perception and imagery respectively.

## Discussion

This study demonstrated that CapsNets are a suitable tool to reach high generalization performance while maintaining a biologically plausible architecture. In previous work, ANNs lacked invariance towards unseen modifications. Our results demonstrated that the proposed network architecture is able to generalize towards unseen locations, sizes, and rotations. The fact that identifiable reconstructions were elicited when occluded stimuli were used as samples indicates that the network also uses information from surrounding (contextual) information in its feedback pathway. The implemented decoder yielded good reconstructions while aligning with fMRI activations. The results of the presented study support the hypothesis that the CapsNet activation patterns show a relevant relation to the brain activation patterns during perception and imagery. Correctly classifiable reconstructions from higher-level visual area activations during imagery indicated that equivariant representations of stimuli are elicited without the necessity of physical stimulation. Since the network that mapped the fMRI activations onto the capsule space was only trained on perception data, the good performance on novel imagery data indicates that perception and imagery share representations in higher-level visual cortex. Lower-level area activations, especially in V1, showed strong similarities to the CapsNet activations. Activation patterns in the encoder related more closely to the perceptual, whereas activation patterns in the decoder related more closely to the imagery fMRI activation patterns. These overlapping representations with neuroimaging activation patterns strongly suggest that implementing a mechanism focusing on a high generalization ability with respect to transformations such as position, size and orientation into a neural network is important to achieve high performance while keeping the architecture biologically plausible.

### Early Visual Area Activations in the CapsNet Overlap with Imaging Activations

On the one hand, perception and imagery could be differentiated in early visual areas: the correlational results revealed that the encoder structure of the CapsNet was more closely related to the perception fMRI activations whereas the decoder structure was more closely related to the imagery fMRI activations. This confirms the idea that perceptual processes rely more strongly on the feedforward pathway whereas imagery processes more strongly involve feedback connections (Koenig-Robert & Pearson, 2021). Building upon these results, it would be an interesting approach to differentiate the cortical layers of the early visual area activations to see whether feedforward and feedback activations can be distinguished. Feedforward activations are assumed to mainly engage middle layers whereas feedback activations are expected to be processed in deep and superficial layers. Therefore, it might be expected that the fMRI perception activations mainly rely upon the middle layers whereas the imagery activations are assumed to rely upon the upper and lower layers. On the other hand, even though there were differences found between the imaging activations alignment with the encoder and the decoder, a strong overlap between perception and imagery depictions were seen in early visual areas, especially in V2. This aligns with the finding that imagery engages early visual areas in a depictive manner, as well as that low-level features are encoded similarly during imagery as they are coded during perception.

To be able to highlight the differences in reconstructions between imagery and perception in more detail, it might be necessary to introduce an adapted pRF mapping for imagery reconstructions. In the presented study, the imagery pRF mapping was calculated based on the perceptual activations elicited while perceiving a checkerboard. It might be that the patches of voxels activated during imagery are more blurry and therefore do not align with the pRF mapping during perception.

### Overlapping Representations in Higher-Level Areas during Perception and Imagery

The overlap in representations at higher-level areas during perception and imagery is consistent with previous findings by Dijkstra et al. (2019). This suggests that the feedforward stream activates higher-level neurons during perception that can then be reactivated to stimulate the feedback stream during imagery. It has been shown that monkey IT includes neurons that fire due to view-specific properties but also a subset of neurons that fire only depending on the specific object presented (Booth & Rolls, 1998; Ito et al., 1995). These findings align with an equivariant representation in higher-level areas in monkey brains. The preserved information in the highest-level area of the CapsNet includes the stimulus category but additionally comprises the relevant information for the reconstruction of the stimulus, differing depending on the specific instantiation of the stimulus.

The utilized fMRI data did not include full extent of IT (since the original study by Senden et al. (2019) focused on early visual cortex), which is a strong limiting factor in comparing the highest layer of the CapsNet and the fRMI activations in high-level visual cortex. Dijkstra et al. (2019) and Koenig-Robert and Pearson (2021) showed that the higher an area is located in the visual ventral stream, the more overlap can be found between perception and imagery activations. Our work showed that even with partial coverage of ventral visual cortex lacking more anterior regions, reconstructions were still possible suggesting that also mid-level visual ventral stream areas exhibit similar representations during perception and imagery. Possibly, reconstructions would have been more accurate if the full ventral visual cortex would have been available.

Furthermore, visual inspection of the reconstructed stimuli from imagery trials did not show the high resolution of Qiao et al. (2018). This might be due to multiple reasons. Firstly, they used activation patterns of voxels throughout the whole brain. The present analysis was restricted to hypothesis-driven higher-level visual brain areas. Moreover, they only reconstructed letters from perception trials. Here, reconstructions from imagery activations were generated. Since imagery is assumed to be a weak form of perception, more noise might be introduced into reconstructions (Pearson, 2019). Thirdly, more trials per category were collected in their study. Therefore, they had more training data, also including more variability between presented stimuli which facilitates the training of the deep neural network that maps brain activation onto capsule activation. Therefore, when testing the proposed network, it should be considered to use a data set with more variability in their stimuli as well as an increased number of trials. The present study serves as proof-of-concept that imagery trials can be reconstructed from mapping of higher-level area activation patterns onto capsule space.

### High Generalization Performance

Our work confirmed that CapsNets provide high generalization performance (Sabour et al., 2017). The results strongly imply that CapsNets are able to classify stimuli independent of their specific qualities, such as size, position and rotation as shown in the presented work. On the other hand, there is evidence that humans perform mental transformations, such as mental rotation, to match representations (Shepard & Metzler, 1971; Koriat & Norman, 1985). Such findings suggest that ‘invariance’ is achieved eventually in multiple ways during recognition and in comparison tasks. An interesting avenue for future work would be to test how the network would behave in situations of ambiguous classifications due to rotations, for instance when “p” and “b” stimuli are supposed to be classified. If similar error patterns as for humans would occur.

A major shortcoming of the present study is the lack of comparison to established networks, such as CorNet-Z (Kubilius et al., 2018), VGG-16 (Simonyan & Zisserman, 2014), or ResNet-50 (He et al., 2016). These networks have shown on certain measures, such as error levels, to replicate human classification performance (Kriegeskorte, 2015) while maintaining a hierarchical structure that has been linked to the human ventral object pathway (Leek et al., 2022). It remains unclear whether these networks can serve as theoretical frameworks for understanding image classification in biological vision (Leek et al., 2022). Adding to the body of knowledge, these networks could be evaluated on the same generalization performance measures as well as compared to the same data set. The lack of an equivalent to feedback connections in these networks will limit such a comparative analysis. Nonetheless, such comparisons might still lead to a better understanding of how different approaches to enhance invariant processing fit to the representations in the ventral visual stream during perception.

Similarities between the fMRI data and the capsule layer activations could be shown but the full extent of the network’s capabilities could not be tested with this limited fMRI data set. The network is specifically trained to generalize to different sizes, locations, and rotations but the data set only included letters presented in the same location, size, and at the same rotation angle. Further research is needed to test whether the differing activation patterns of the network also reflect the different activation patterns for other locations or sizes when human participants perceive stimuli. For instance, it would be interesting to investigate whether the reconstructions of stimuli imagined in different locations are also projected onto different locations.

### Image Comparison Measurements

The generalisation performance was not only measured with classification accuracy and loss calculations but also with the help of correlations, RMSE, Euclidean distance, PSNR, and SSIM. These measures are similarity measures that compare two images with each other and they all rely on low-level feature similarity (Rakhimberdina et al., 2021). All of these measures were used to gain a broader perspective on similarity than using only spatial correlation measure as in Senden et al. (2019).

Interestingly, when using correlations as a measurement, high similarities were found but the SSIM, as well as the PSNR, indicated low similarities. This divergence between the results might be explained by the different shortcomings of each of them. Correlations are often used but lack sensitivity to changes in the edge intensity and edge alignment (Beliy et al., 2019). MSE is also a regularly applied measure for comparisons, but it shows poor correspondence to human visual perception. MSE calculations are independent of the spatial relationships between image pixels and do not give differing weights to each of the pixel (Wang et al., 2004). The same applies to RMSE which was used as standardization of MSE in this study. The same problem also arises with Euclidean distance which is defined as a summation of the squared pixel-wise intensity differences. Therefore, even small deformations can result in large distance values (Wang et al., 2005). PSNR and MSE show low discrimination power for structural content as demonstrated by multiple degradations applied to the same image yielding the same value of the measurements (Hore & Ziou, 2010). This problem arises since MSE and PSNR are giving absolute errors. On the other hand, SSIM was proposed as being a similarity measure that captures human visual perception. It calculates a weighted combination of three comparative measures which are luminance, contrast, and structure (Rakhimberdina et al., 2021). Even though this measure is correlated with human quality perception, it still compares poorly to some human perceptual characteristics (Zhang et al., 2018).

As a potential improvement to the present study, higher-level perceptual similarities should be taken into account when comparing samples with reconstructions. Therefore, the PSM (Perceptual Similarity Metric) is proposed and should be evaluated in further research on CapsNets. This metric uses a CNN to extract hierarchical multilayer features of input images. These are then further compared across layers using a distance metric. Often, AlexNet is used in that context to extract the hierarchical features (Rakhimberdina et al., 2021).

### Biological and Psychological Plausibility

In line with Svanera et al. (2021), we observed that a network with an encoder and decoder can utilize surrounding information to reconstruct occluded stimuli. Differently from their study, this network was not particularly trained for reconstructing occluded images but showed this behavior following training that optimized classification and reconstruction. Our network and training regime are in this regard more biologically plausible (Smith and Muckli, 2010).

We also tested the network’s performance with the original decoder from Sabour et al. (2017) but results were subpar and the network classified images randomly. This was to be expected since the stimuli showed increased variability compared to MNIST images. Nonetheless, just increasing the size of the layers or the number of layers did not lead to better results. Instead, the main architecture from the feedforward network was reversed which also relates to the hypothesized architecture in the brain, using a feedback stream that leads through the same areas as the feedforward stream (Dijkstra et al., 2019).

Even though a decoder has been implemented in the network, one might argue that these so-called feedback connections are not informative of the human brain since they lack influence on layers earlier in the visual stream. For instance, it has been shown that neurons in early visual areas show target-enhancement, particularly observed after longer latencies which implies that not the on-discharge is relevant for the activation (Hon et al., 2009). Instead, feedback about target presence was assumed to influence early visual areas. Besides a different implementation of feedback connections, also adding lateral connections might make ANNs more biologically plausible (Kubilius et al., 2018).

A further improvement of this visual ventral stream model might be the introduction of an attention mechanism (Goebel, 1992). Humans are well able to outperform ANNs once the presented classification tasks become more challenging because they are able to direct their attention towards relevant parts of a scene while ignoring distracting information (Peters et al., 2012). Different attempts to implement attention mechanisms with capsule networks were made and showed promising results in object classification tasks (Huang & Zhou, 2020; Mazzia et al., 2021; Pucci et al., 2021). Implementing an attention mechanism in the proposed network might lead to improved explanation of human activation patterns.

Another example of psychological constraints that might be tested to claim similarities to human performance is related to feedforward deep neural networks classifying stimuli depending on local shapes and texture instead of global shapes (Baker et al., 2020). AlexNet, VGG-19, and ResNet-50 were trained to classify squares and circles. For testing, squares comprised of curved elements and circles comprised of squared shapes were presented. For classification, the networks relied upon local contour features rather than global shapes. They argued that global features would dominate human vision whereas ANNs only represent how local contour fragments spatially relate to each other when forming global shapes. Since the routing-by-agreement algorithm strongly relies on the feature relations of an object, this paradigm might be a promising additional test to demonstrate the generalization ability of our network and CapsNets in general.

### Interpretability

Current machine learning approaches modelling human brain activations frequently do not provide an understanding of the underlying neural processes or how they might contribute to the outcome of the network (Fellous et al., 2019). Additionally, high-performing methods tend to be the least explainable thereby providing little additional understanding of mechanisms in the human brain. In recent years the interest in explainable artificial intelligence rose (Angelov et al., 2021). Since Sabour et al. (2017) claimed a linear relationship between the capsule layer activations and the reconstructions, it was thought that the presented network might offer a higher degree of explainability. When testing this hypothesis, it became clear that even though this network contains some form of equivariant information at the capsule layer, this representation seems to contain an abstract combination of features which could not be easily interpreted.

## Conclusion

The presented CapsNet provides high generalization ability with respect to object transformations as well as alignment with fMRI activation patterns in lower-level as well as higher-level areas in the visual ventral stream. These promising results suggest that the inherent high generalization ability of CapsNets might be relevant in implementing biological vision. The decoder structure which implements feedback connections inverting the CapsNet architecture seem to be an improvement compared to networks relying only on feedforward recognition. Additionally, high-level areas indicated coherent activation patterns during perception and imagery. This finding demonstrates an overlapping representation during the two different tasks. Further comparisons to other established networks are needed to distinguish whether the high generalization performance is a necessary characteristic for improved modelling of the ventral visual stream.

## Appendix

All used code and data from the presented study can be found on GitHub at: https://github.com/FKlepel/Thesis_CapsNet_Perception-Imagery

